## Supplemental figures for "A humanized yeast model for studying TRAPP complex mutations; proof-of-concept using variants from an individual with a *TRAPPC1*-associated neurodevelopmental syndrome"

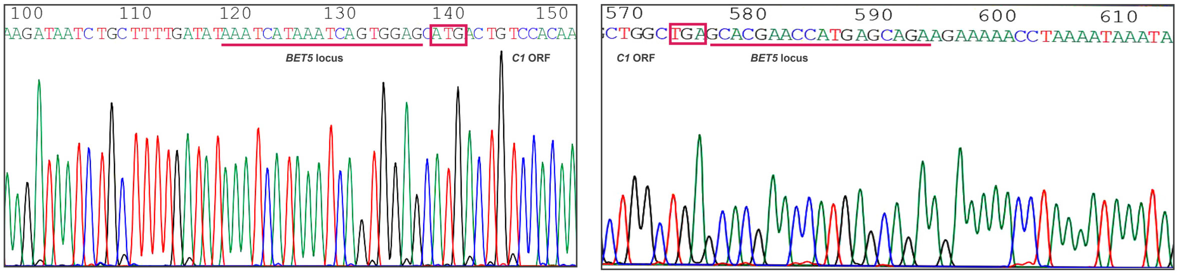


Humanized yeast with *C1*

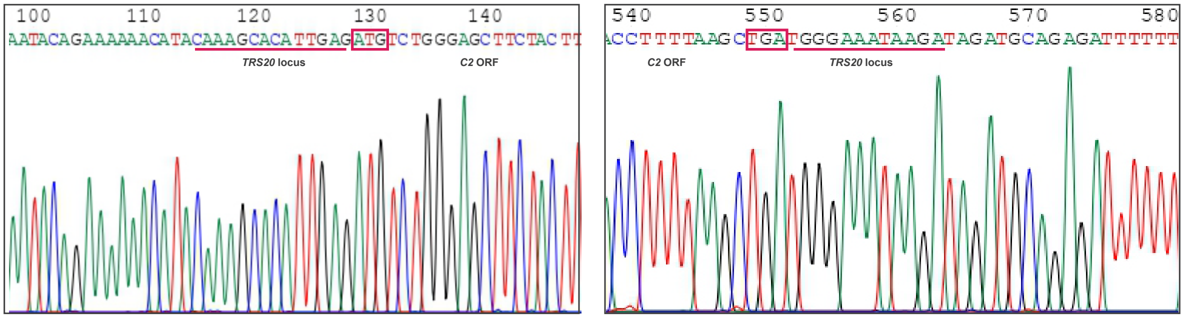


Humanized yeast with *C2*

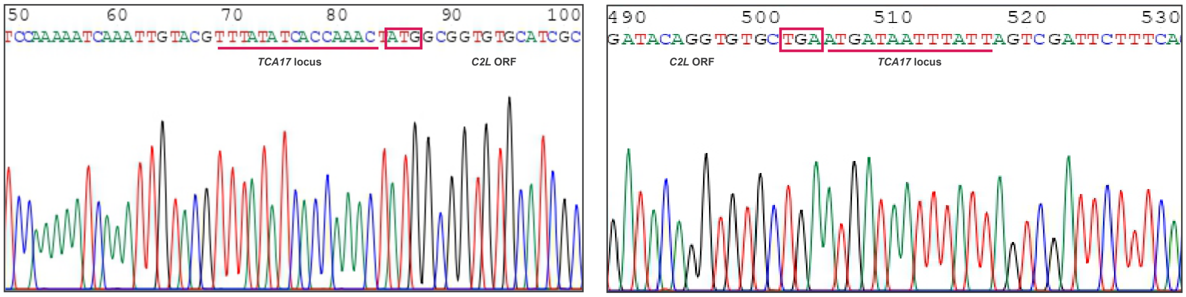


Humanized yeast with *C2L*

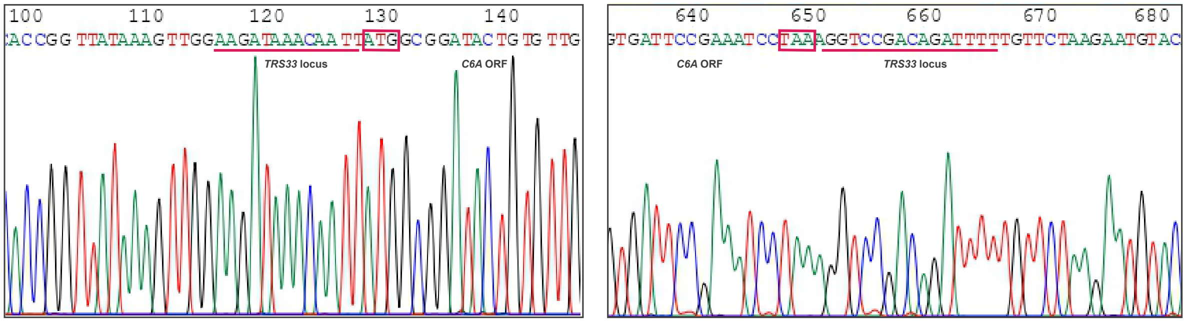


Humanized yeast with *C6A*

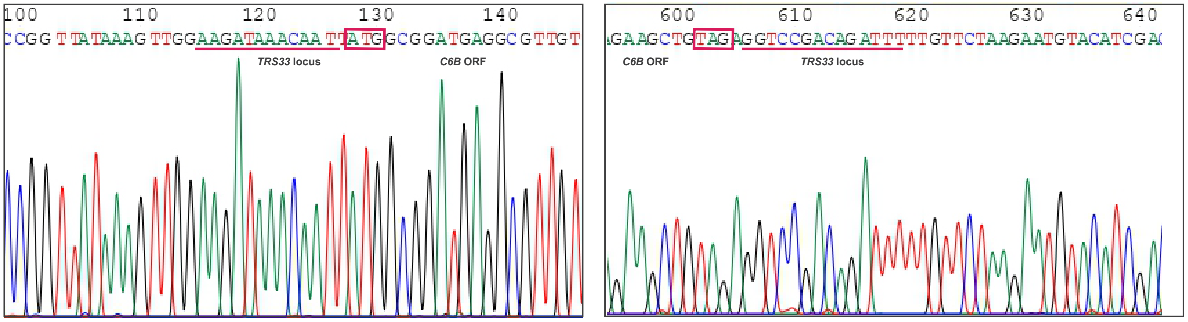


Humanized yeast with *C6B*

### FIGURE S1

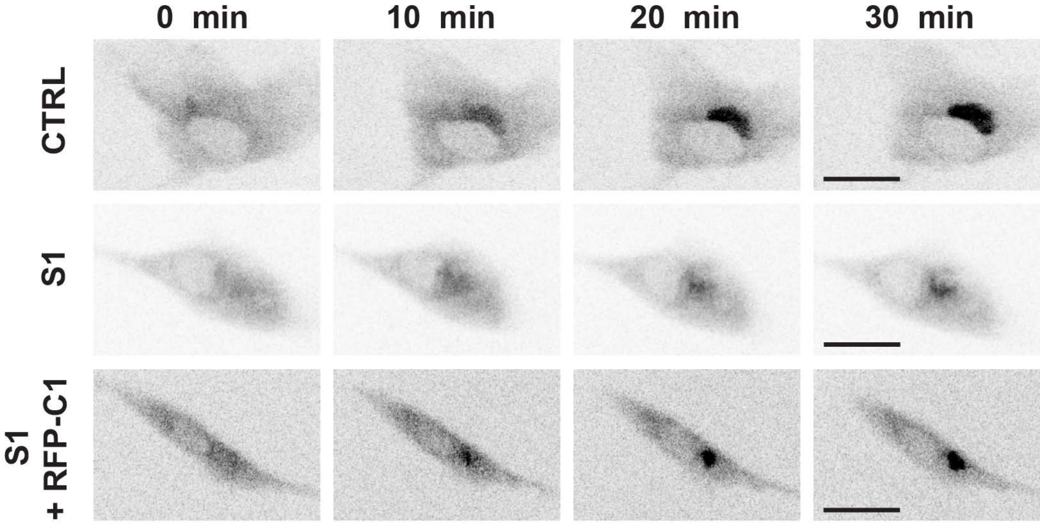


### FIGURE S2
